# Linking coleopteran diversity with agricultural management of maize agroecosystems in Oaxaca, Mexico

**DOI:** 10.1101/2020.01.07.897744

**Authors:** Cecilia González González, Tania Lara García, Lev Jardón-Barbolla, Mariana Benítez

## Abstract

Biodiversity is known to be influenced by agricultural practices in many ways. However, it is necessary to understand how this relation takes place in particular agroecosystems, sociocultural contexts and for specific biological groups. Also, in order to systematically study and track how biodiversity responds or changes with agricultural practices, it is necessary to find groups that can be used as practical indicators. We conduct a study of beetle (Coleoptera) diversity in maize-based agricultural plots with heterogeneous management practices in the Central Valleys of Oaxaca, Mexico. First, we use a mixture of local knowledge and multivariate statistics to group the plots into two broad and contrasting management categories (traditional vs. industrialized). Then, we present an analysis of Coleopteran diversity for each category, showing higher levels across different diversity indexes for the traditional plots. Also, our results let us postulate the Curculionidae family as an indicator of both management type and overall Coleopteran diversity in the agricultural lands of the study site. We discuss our results in terms of the agricultural matrix quality and its role in joint productive and biodiversity conservation strategies.

## Introduction

Agriculture is a process in constant evolution that depends on the political, economic and cultural processes of the places where it develops (Vandermeer 2011). As such, it is a diverse phenomenon composed by a large variety of productive practices that employ different tools, techniques, species, both vertical and horizontal arrangements, and very importantly, knowledge and human labor. These practices are not assembled at random, but occur as sets of land management activities that are interdependent, adapted to each other and that function as a system with specific goals (Andow & Hidaka 1989; Vandermeer 2011). Here we will use the term “management type” in order to name these sets of interdependent productive practices with which human labor produces from the land.

Starting with the industrial era and ever since, a homogenizing tendency has changed the ways in which agriculture is done (Perfecto et al. 2009). Technologically, the industrialization of agriculture has implied a gradual transition from systems with a high planned biodiversity (those species that the peasants decide to actively maintain or grow in the agroecosystem) and a low or null import of inputs, towards systems with a low planned biodiversity and a high dependence on imported inputs that in some measure seek to replace the functions of the removed elements of biodiversity and compensate the nutrient depletion that comes from an exhaustive use of resources (Perfecto et al. 2009). In Mexico, we see the full expression of this process in the large farms of the maize belt at the northern part of the country. Nevertheless, more to the South, where this study was done, one finds a very heterogeneous mixture of practices that are carried out by small producers in plots that are usually no bigger than 5 ha (INEGI 2007).

The homogenizing tendency described above has had profound effects on biodiversity, as the Secretariat of the Convention on Biological Diversity (2014) has stated: 70 % of the biodiversity loss projected to the near future is associated to the agro-alimentary system. However, there is not only one type of agriculture, but a vast diversity of management types that have markedly different effects on ecosystems; and some are capable of maintaining or even incrementing biodiversity at a landscape level (Perfecto 2003). The classic conservation approach that understands agriculture as an absolute antagonist to biodiversity has in mind an industrialized agricultural management, but we need to overcome this simplistic view and assess agriculture as the heterogeneous process it is, in order to build alternatives that can feed us and conserve biodiversity at the same time. This is particularly important for agricultural systems near or within biodiversity hot spots that are associated to the production of culturally meaningful foods (Bellon et al. 2018).

Indeed, different management types have different effects not only on the planned biodiversity but also on associated diversity, that is, those species that are not actively maintained by peasants but that are capable to establish temporally or permanently at agricultural lands, or to migrate through them (Perfecto et al. 2009). For example, traditional coffee farms have been found to hold a diversity of canopy insects, birds, mammals, plants and other organisms as high as that from conserved tropical forests in South America (Perfecto et al. 1997; Perfecto & Ambrecht 2003; Perfecto et al. 2007). In contrast, in industrialized coffee farms, the implied absence of tree canopy has led to a drastic decrease in these same taxa (Perfecto et al. 1996; Greenberg 1997; Philpott et al. 2008). The same tendencies have been found in cacao agroecosystems (Perfecto et al. 2009), rice (Bambaradeniya 2000) and maize (Finegan & Nasi 2004).

In addition to being biodiversity repositories, agricultural lands also have a part on metacommunity processes, as they are often in the way of migration across conserved patches of forest. In this manner, an agricultural management that opts for crop diversity, preventive plague control, soil conservation and a heterogeneous spatial structure can foster the connectivity and permanence of functional metacommunities (Perfecto et al. 2009, Tscharntke et al. 2012; Kremen 2018).

Once that we recognize the importance of discerning different management types, we can ask what kind of indicators are useful for their practical identification and monitoring. Because organisms’ response to perturbation differs greatly among taxa, there is not one single organism that can serve as an indicator, but it is advisable to combine indicators from different groups as plants, vertebrates and invertebrates (Schulze et al. 2004). Nevertheless, the order Coleoptera has been found to correlate with other vertebrate and invertebrate taxa (Pearson & Cassola 1992; Holland 2002; Ohsawa 2010). In addition to serving as an indicator of other species, beetle diversity also has been found to correlate positively with environmental variables like soil quality and habitat complexity (Lassau et al. 2005; Campanelli & Canali 2012), aspects that in turn relate directly to management type in agroecosystems.

The reasons underling the potential of the order Coleoptera as a good source of biological indicators are plenty: they are among the most abundant beings in the world (they form almost 40 % of insects and 25 % of animals), they are cosmopolitan, they define a great variety of niches and they maintain several ecological interactions making them important matter and energy flux regulators. Moreover, they are relatively easy to sample and have a great variety of life strategies involving processes with landscape-level implications like reproduction, dispersion and colonization (Hunt et al. 2007; Bouchard et al. 2011). The revisions by Kromp (1999) and Holland and Luff (2000), and the work by Burgio et al. (2015) are good examples of the family Carabidae being strongly influenced by different agricultural management practices, and the work by Brooks et al. (2012) use this same family as a biodiversity indicator. Yet, the usefulness of beetles as indicators is highly context-specific, so results should not be extrapolated to different geographic areas, rather, it is recommendable to study which families are useful in each site (Jonsson & Jonsell 1999).

For these reasons, we sampled the beetle community in order to find differences in its diversity for contrasting management types, and then we searched for possible indicators of both management and biodiversity. We worked in the Central Valleys of Oaxaca, Mexico, a region recognized for its extraordinary biological, agronomic and cultural diversity (Mora 2017).

## Methods

### Study Area

This study was conducted at Villa de Zaachila (Zaachila hereafter), in the Central Valleys of Oaxaca, Mexico. It is a semi-urban population located 17 km southeast of the state’s capital. The most represented land use types are agriculture, which covers 48 % of its total area, secondary forest with 23 % and urban zones with 19 % (Urrutia at al. 2019). According to government data, the municipality is composed of 1,669 ha which are distributed among 1,521 peasants (heads of family), which is to say every peasant family has an average of 1 ha of land to work with. This small-scale agricultural scheme combines with the fact that 90 % of the lands are rainfed while 10 % are under irrigation; 66 % of the peasants use both machine and animal traction, while 27 % employ only tractors and 7 % use only animals; 41 % apply industrial fertilizers; 4 % use some type of pesticide and only 1 % use hybrid seeds (INEGI 2007). Thus, there is a wide heterogeneity of agricultural practices in the area combined at a rather fine scale.

The history of landscape management of Zaachila begins with the Zapotec peoples, about 3500 years ago. Throughout its history, the municipality has had different management and land holding systems (Ruiz Medrano, 2011). Today, agricultural plots are mostly managed by peasants for family or local consumption. In contrast with most agricultural sites in other countries, many of these plots represent a state-given usufruct land, the so-called *ejidos*, in which variable degrees of collective management occurs (INEGI 2010; Mora 2017). Historically, Zaachila has been an important point for regional commerce, as its traditional market has existed since the time of the Zapotecs, and even now it gathers farmers and peasants from all the surrounding villages. Zaachila is an important trade center, since its weekly market has attracted peasants selling their products from all round the Central Valleys region since prehispanic times (Aguilar & Huebe 1979). Crops such as maize, beans, peanuts, alfalfa and walnuts are sold here every Thursday, making up an essential part of the local peasant economy (Mora 2017).

### Plot selection

We looked for plots with contrasting management types, which we will call *traditional* and *industrialized*. For this we used the methodology developed by Álvarez et al. (2014), which seeks to group heterogeneous plots based upon local key informers (mainly, the peasants and their families) and multivariate statistics. The method’s goal is to obtain a reproducible categorization of the management types in a certain location, which serves both the needs of the investigation and the perceptions of those involved.

During september 2016 we worked with a group of peasants with whom we had built a trusting relationship since two years earlier due to previous activities in the area (Benítez & Jardón-Barbolla 2015; Mora 2017; Urrutia et al. 2019). Together, we set to find plots belonging to two qualitatively different management types. First, a *traditional* type, characterized by a low use of commercial inputs and the planting of diverse, local seeds. Even though the exact combination of agricultural practices in each plot actually varied, we postulated that by sharing those two features they would constitute an ecologically similar category, so validity of this prior classification was tested through the information we got. Second, we looked for the industrialized type, which would be formed by plots with a high use of machinery, agrochemicals and hybrid seeds. With this in mind and according to their knowledge of local practices, the local peasants led us to 16 plots, half belonging to each type, and distributed along the North, South, East and West parts of the locality, having plot pairs in each zone.

In order to test the hypothesized categorization, we selected 22 quantifiable management variables with which to describe each plot, such as number of grown crops, presence of trees inside the plot, use of organic or non-organic inputs, use of machinery, irrigation, and so forth. The complete list of variables, definitions and data type can be read in in the supplementary material (TableS2).

### Interviews

We conducted semi-structured interviews with the owners of each plot in order to collect information on the 22 variables mentioned above. These were held in the peasants’ homes or in their plots, and were recorded in audio and paper via summarized questionnaires. Some of the variables were also verified by sight in the plots, such as the presence of trees or herbs. Prior to this process, an informative letter was given to each of the participants, in which we explained to each family the exact purposes of our research, as well as our academic affiliation. All participants gave participants informed consent to participate in the study.

Data was later translated into a quantitative table, with two variables (number of crops and number of crop varieties) recorded as absolute richness values and the rest of the variables as binary presence/absence values. We are aware of the loss of information that this process implied but we hold it necessary for having a comparable set of data for all plots and to proceed as suggested by Álvarez and collaborators (2014). This table can also be found in the supplementary material (Table S3).

### Testing the plot categorization

We created a correlogram of Spearman coefficients in order to look for relationships among management practices. Then, due to the type of variables in our study, we used a Factor Analysis of Mixed Data (FAMD) for grouping the plots according to their management practices. All tests were done using the *FactoMineR* and *corrplot* packages from the coding language *R 1.1.383* (R Core Team 2014). All scripts used for this investigation are publicly available at: https://github.com/laparcela/Coleoptera. The groups resulting from this categorization were later used as treatments to compare beetle diversity.

### Beetle sampling and identification

Plot extension was around 50 × 200 m, and we established five quadrants measuring 1.5 × 1.5 m in each one. We first established a random point in each of the long sides of the plot and drew an imaginary line connecting them. At this line’s center, we established the first sampling quadrant. Then we established another two quadrants halfway between this center and each of the line’s extremes. Afterwards, a second imaginary line was drawn perpendicularly to the first line, crossing its center. The fourth and fifth quadrants were established at the extremes of this second line. Following this procedure, we obtained five quadrants in each plot, three in their interior and two at their borders (A diagram can be found in the supplementary Figure S1).

During September 2016, towards the end of the rainy season and when crops were ready for harvesting, all plots were sampled through the use of sweep nets in the five quadrants. Two plots were sampled each day, starting at 7:00 am and until approximately 11:00 am. This was kept constant in order to avoid extra noise from a day’s normal temperature variation. Collected specimens were fixed with 70% ethanol and aggregated as a single sample for each plot and then taken to a laboratory for identification. Individuals were identified to family level using identification keys (White 1983; Tripplehorn & Johnson 2005) and to morph level afterwards. Diversity among the industrialized and the traditional plots was later compared using one-way ANOVAs, Renyi diversity analysis (using the *vegan* and *BiodiversitiR* packages from the coding language *R 1.1.383* (R Core Team 2014)) and direct quantifications at several taxonomic levels.

## Results

### Plot categorization

The correlogram showed relations among sets of management practices, as expected from the theoretical postulation of management types (Vandermeer 2011; Álvarez et al. 2014). In Figure 1 we present Spearman coefficients among the 22 chosen variables. We found a positive correlation (groups of blue circles below) between practices commonly associated to a traditional management, such as managed border, use of manure, animal traction, number of crops, quelites (non-planned herbs that are frequently useful to peasant families in different ways), number of varieties, presence of trees and use of organic fertilizers. Likewise, there is a strong positive correlation between practices associated to an industrialized management, such as use or industrial fertilizers, herbicides and pesticides, irrigation and monoculture. We also found negative correlations (groups of red circles) between irrigation and fallow time, and subsistence agriculture and monoculture.

**Figure 1.**
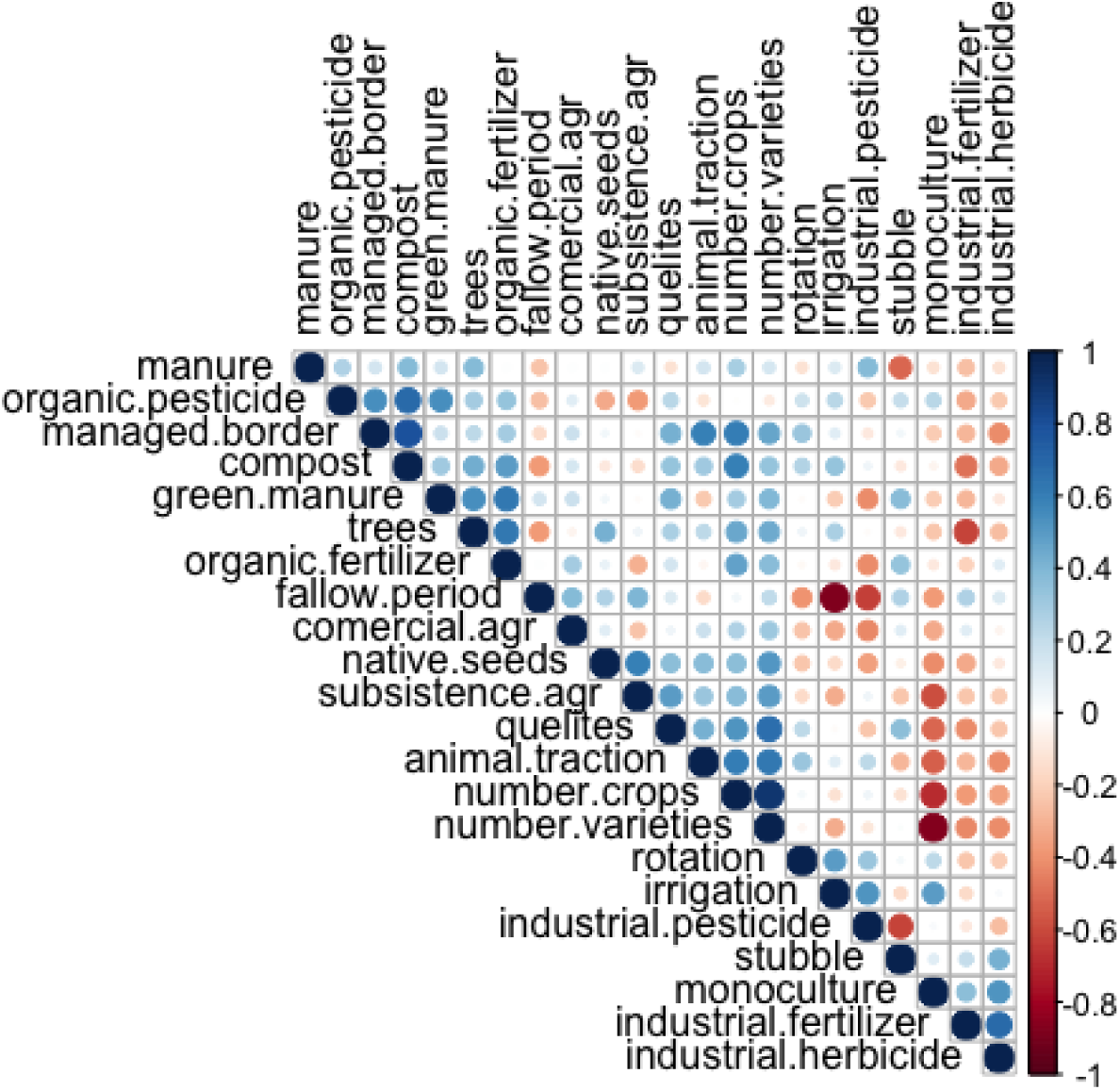
Spearman correlation coefficients between management practices.

The FAMD we conducted explained 46.2 % of the total variance in its two first dimensions (27.4 % and 18.8 % respectively). In figure 2.b is the variables factor map, where we show that the most correlated variables with the first dimension were the number of varieties within crops (0.896 correlation), number of crops (0.872 correlation), and the monoculture planting scheme (−0.722 correlation); while the second dimension was most correlated with the presence of a fallow period (0.9 correlation), irrigation (−0.84 correlation) and use of industrial pesticides (−0.62 correlation) (see TableS2 for a more detailed definition of each variable). Moreover, all the variables with a positive correlation to the first dimension were practices associated to a traditional management (Altieri et al. 1997): use of local seeds, crop rotation, fallow period, animal traction, presence of quelites, managed border, presence of trees, use of green manure, compost regular manure and organic inputs. On the other hand, the variables with a negative correlation were all associated to an industrial model of agriculture (Perfecto et al. 2009): a monoculture scheme, irrigation and use of industrial inputs (fertilizers, herbicides and pesticides).

**Figure 2.**
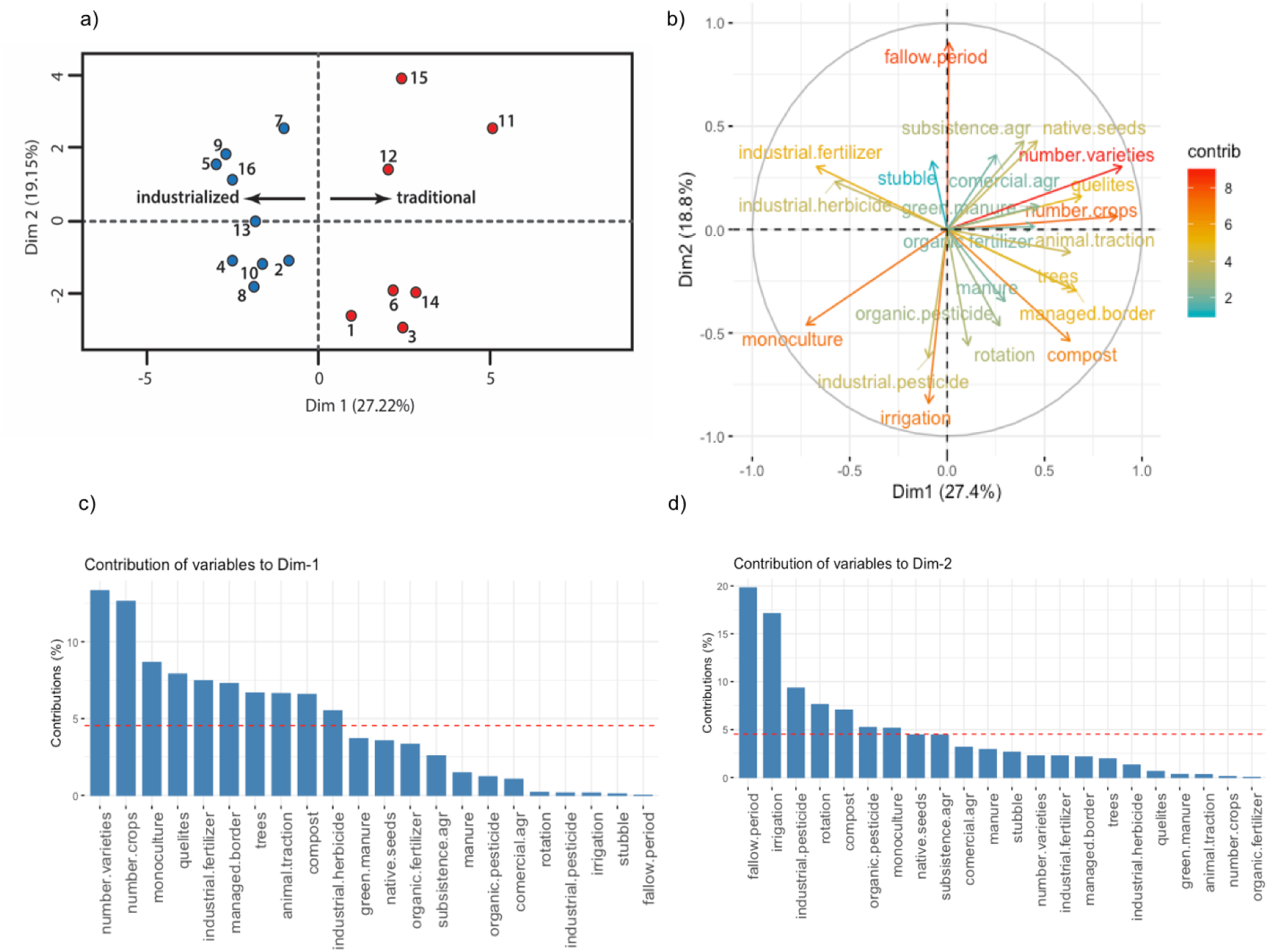
FAMD analysis. a) FAMD output for the 16 plots under study, plots on the left side correspond to the industrial management type and plots to the right correspond to the traditional management; b) variables factor map, colors are given by each variable’s contribution to the first and second dimensions; c) contribution of variables to the first dimension; d) contribution of variables to the second dimension. The reference line in c) and d) corresponds to the expected value if the contributions were uniform.

The first dimension of the FAMD depicts a management gradient that goes from the most industrialized on its negative side to the most traditional on its positive side (Figure 2.a). Based upon this, we separate the plots falling to the right and left side of the first dimension’s origin, thus discretizing the management gradient into two separate categories (industrial and traditional). By doing this, we found that 15 out of 16 plots fell in the expected side from the hypothesized management category. Thus, we sampled and compared coleopteran diversity among these two groups, leaving aside the one plot that did not match the original hypothesis. Note that the vertical axis was not used to further divide the plots in more groups, nevertheless it is worth saying that the variables with higher coefficients in this dimension were fallow period with a positive sign, and irrigation and pesticide use with a negative sign.

### Coleopteran diversity

In Figure 3 we present an overview of the sampled diversity of families in the order Coleoptera. In total, we found 1168 individuals belonging to 25 families, which we then identified as 80 different morphs. Because the sampling was done with sweep nets, we mainly captured beetles that are active in the above-ground part of the agroecosystem, mainly on stems, leaves, flowers, fruits or sometimes flying in the air. For example, Chrysomelidae, Cantharidae, Curculionidae and Coccinellidae are commonly found on leaves and stems; while Cleridae and Phalacridae are more abundant on flowers and trees.

**Figure 3.**
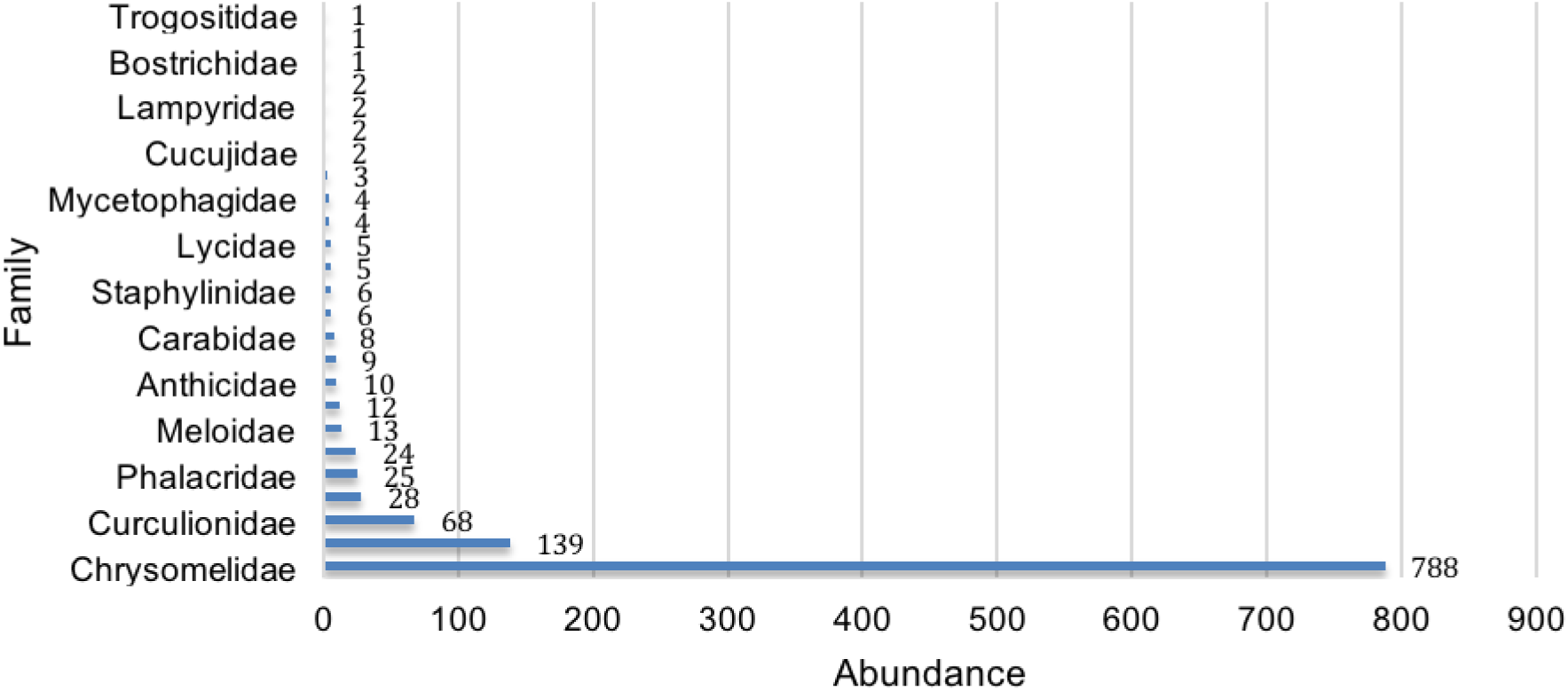
Total abundance of every beetle family collected.

In order to compare the diversity found among plots in our two management categories, we first performed one-way ANOVA tests (Figure 4). Overall coleopteran abundance did not show significant differences among plots from the industrialized and the traditional groups, but family richness was significantly (*p*=0.0265) higher in the traditional plots. Exploring a further taxonomic subdivision, we found that morph richness was also significantly (*p*=0.0203) higher in these plots.

**Figure 4.**
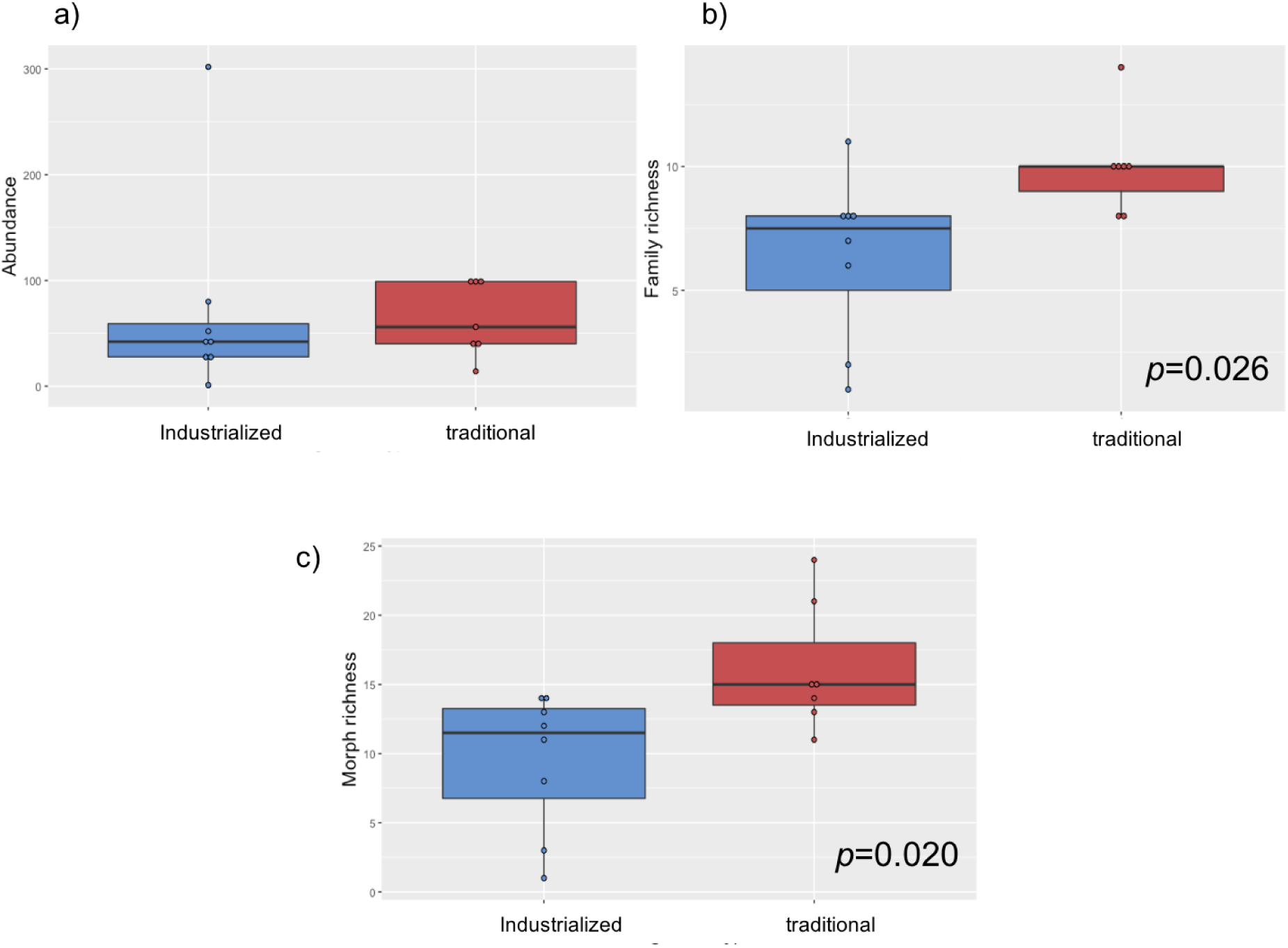
Boxplot and ANOVA tests for a) individual abundances, b) family richness and c) morph richness. The median is represented by the horizontal line inside the boxes, lower and upper hinges correspond to the first and third quartiles, whiskers extend to the largest value no further than 1.5 times the inter-quartile range, data points are represented by circles and data beyond the whiskers are outliers. The ANOVA assumptions of equality of variances and residual normality were covered by the data: Bartlett test *p*=0.1551 and Shapiro test *p*=0.4861 for family richness data and Bartlett test *p*=0.6805 and Shapiro test *p*=0.9387 for morph richness.

Following the ANOVA tests, which were performed using each plot as a separate sample, we grouped all plots belonging to the industrialized group in one, and all plots belonging to the traditional group in another sample. In Figure 5.a we show a visualization of morph richness inside each family in these two compound samples.

**Figure 5.**
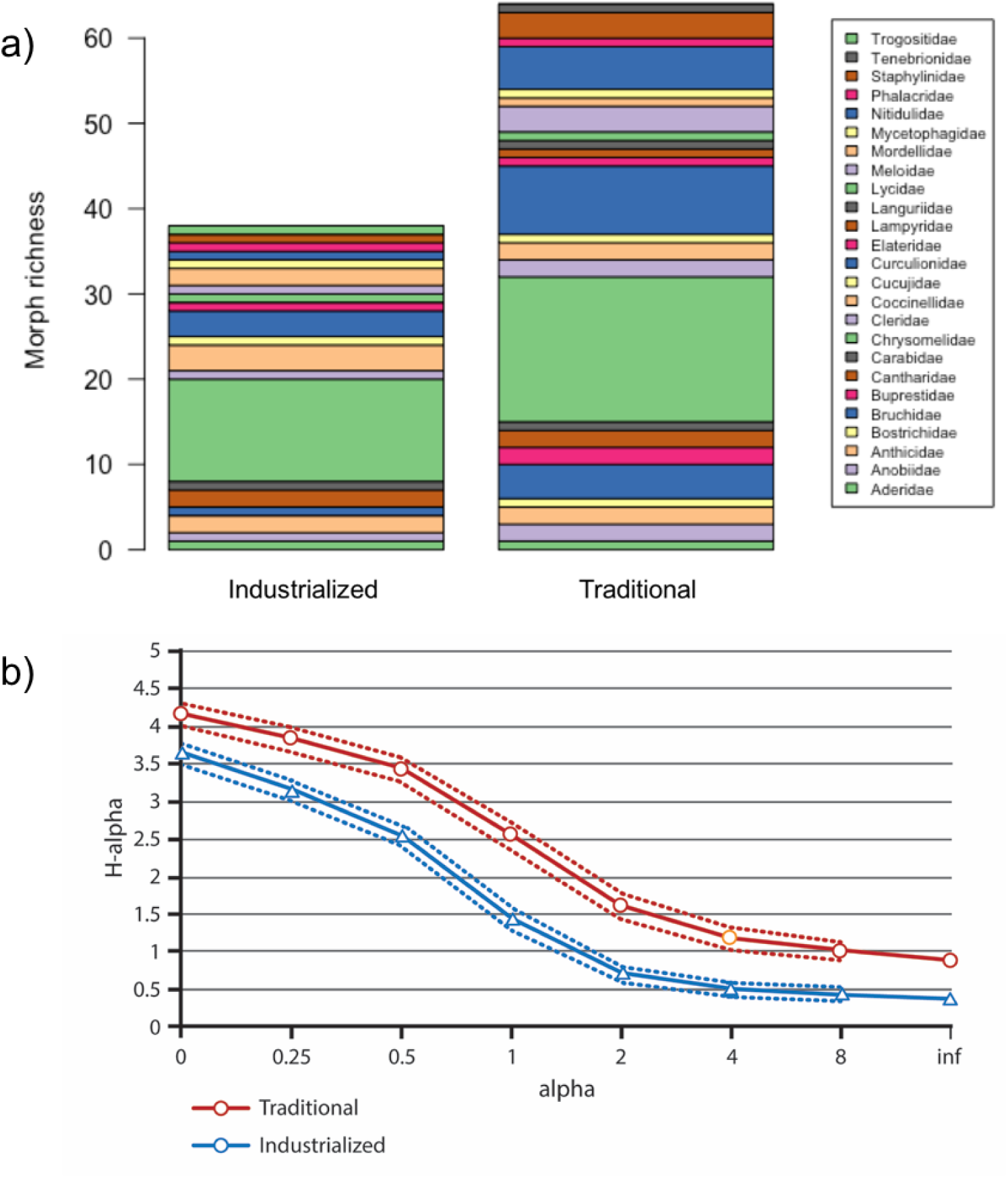
Comparison of the industrialized vs. traditional compound samples. a) Morph richness is strikingly higher for most of the families in the traditional group; b) Renyi diversity is consistently higher in the traditional group.

Then we calculated Renyi diversity for each group and found the traditional one to be significantly more diverse across all the Renyi’s spectre of indexes (confidence intervals at 95 %) (Figure 5.b). Renyi’s analysis relates in a graphical way many widespread ecological indexes, which are located along the horizontal axis while the vertical axis shows the corresponding value for Renyi’s index (H-alpha)^1^.

### Curculioidae as indicator of management type and diversity

Given our results, we then looked for beetle families that could be useful indicators of the agricultural management type. First we noted that there was one family that was exclusive to the industrialized management: Trogositidae. Likewise, there were four families exclusively found in the traditional plots: Bostrichidae, Buprestidae, Languriidae and Tenebrionidae. Nevertheless, these were regarded as non useful because their abundances were extremely low (the most abundant family was Tenebrionidae with three individuals). Thus, we turned our attention to a family that was not exclusive of a particular type of management, but showed a different response to the management categories: Curculionidae (Figure 6.a).

**Figure 6.**
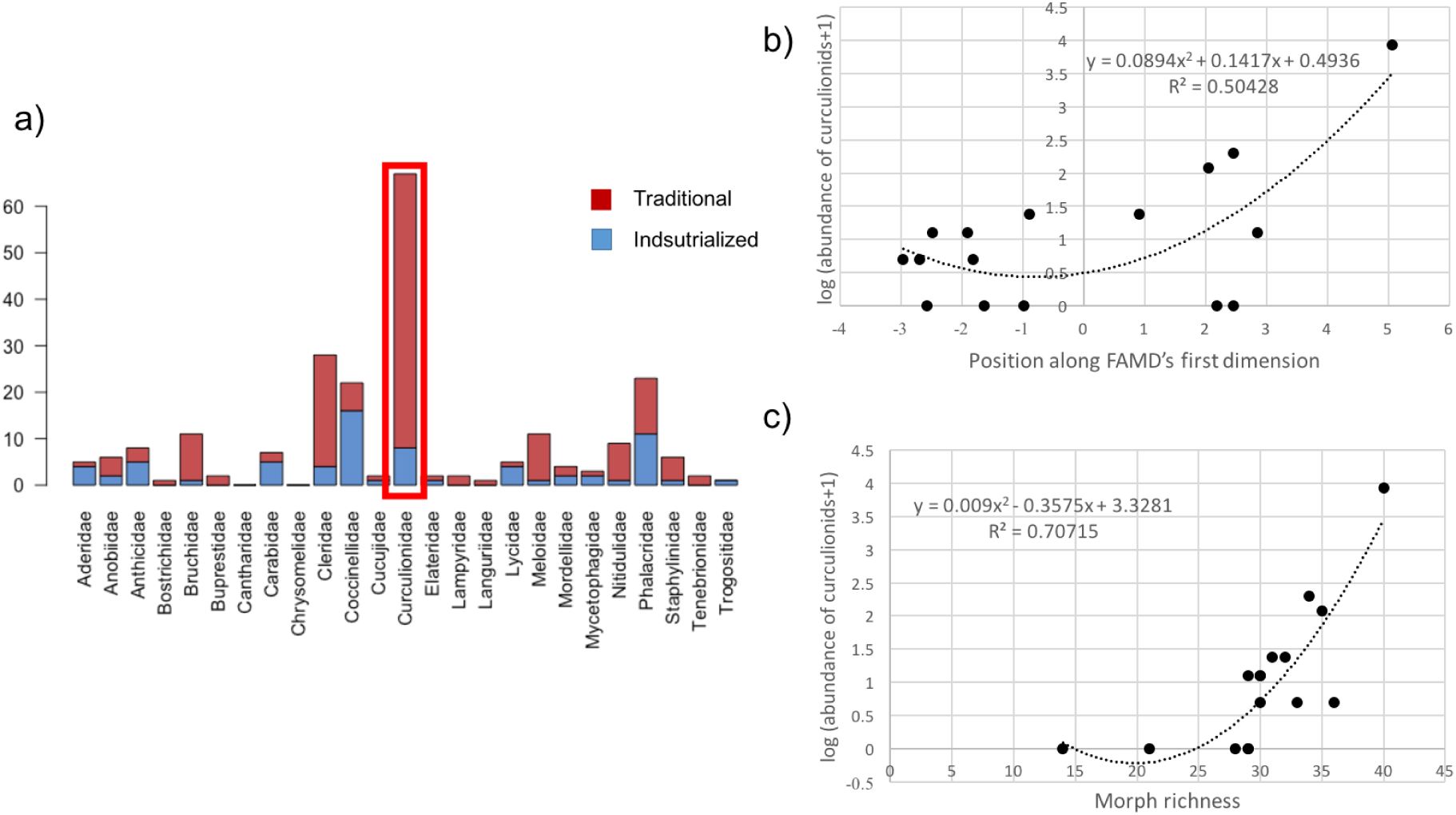
Curculionids as indicators. a) abundance of each family in traditional and industrialized plots, b) polynomial regression of curculionid abundance against management gradient, c) polynomial regression of curculionid abundance against richness of beetle morphs.

In order to test the potential of curculionids as indicators, we analyzed the relation between their abundance and plot management (Figure 6.b). For this, we adjusted a second degree polynomial to each plot’s curculionid abundance (we took the natural logarithm of this measure in order to comply with the analysis assumptions) against its position along the management gradient (the coordinate each plot occupied in the horizontal axis of Figure 2.a. We found a positive relation between these variables (R=0.50428, p<0.05), suggesting that plots with a more traditional management tend to have a higher abundance of curculionids.

Finally, we analyzed the relation between curculionid abundance and the total richness of beetles in each plot (Figure 6.c). We first took richness at a family level but found no significant results, nevertheless, the same comparison at a morph level did show a significant positive relation between the two variables (R=0.70715, p<0.05).

## Discussion

In the first part of our study, we used the method by Álvarez et al. (2014) to define two broad management categories based upon a wide series of agricultural practices. Based on the coefficients of the statistical approach, we found two variables to be most important for the organization of plots into these categories: total number of crops and total number of varieties grown. In a practical sense, this result is interesting because these variables can be measured with relative simplicity by academics, technicians or peasants, and they seem to work as ‘umbrella’ variables for many other practices associated to agricultural management, making them useful guidelines for future work that seeks to consider this matter. From an ecological point of view, crop diversity is involved in many processes both below or above ground. For example, plant diversity in associated to structural diversity (Del Río et al. 2003) and therefore, to the set of niches available for the biota (feeding, refuge, reproduction spots, etc.) inhabiting or passing through the plots.

Another remarkable aspect is the difference in the grouping of the industrialized and the traditional plots. As seen in Figure 3, the industrialized plots are much closer to each other than the traditional plots. This means that even though both categories have a high internal variation, the industrialized plots are more similar to each other than the traditional ones, reflecting the tendency to homogenize that is characteristic of industrialized agriculture in contrast with the heterogeneity and context-specificity of traditional agriculture (Gliessman 2015). The high variability found inside the traditional group of plots is surprising given the small scale of this work, which shows the heterogeneity that can be found even among traditional peasants belonging to the same community. This should compel towards a reflection on the *traditional* management not to be seen as “fixed” or archaic, as there’s relevant variation in the practices that peasants adopt through time and space.

As for coleopteran diversity, the traditional plots showed higher values at both family and morph level, regardless of the diversity measure used and in spite of the fact that many of the industrialized plots were close to them (which could have otherwise brought diversity to industrialized plots). This indicates that coleopterans are significantly sensible to agricultural management even at small scales, a fact worth noting given that local communities are influenced by their surroundings and that their diversity depends on the species pool at a broad scale (Duelli et al. 1999) and on many landscape variables at a small and medium scale (Gabriel et al. 2010). Moreover, the variability inside each management category was also large, as we have discussed above, which makes significant inter-category differences more striking. In addition to having a higher diversity of beetles in general, the traditional plots had more exclusive species, which points to their importance as reservoirs of rare species.

Regarding the family Curculionidae, we found that the abundance of this family increases as management tends to be more traditional and that it also reflects the diversity of coleopterans in general. Nevertheless, this should be taken with caution because it was the case for most, but not all the sampled plots. This said, curculionids can be useful indicators because their morphology is easily recognizable (mainly because of the anterior projection in their head) and in fact it is safe to say that all peasants in Mexico are well familiarized with them. The family is abundant in most of the world, and more markedly in the tropics. They mainly feed on vegetal tissue, though they can also eat lichens, algae and fungi (Morrone 2014). According to Zimmerman (1994), all angiosperms are probably consumed by at least one curculionid species. However, they are not just predators, for various plants depend on curculionids for pollination (Morrone 2014). This family has been used as an indicator of coleopteran diversity before (see Ohsawa 2010), but we have no knowledge of it being used in Mexico or in maize-based agroecosystems. Because of its cosmopolitan nature, it would probably be found to be a good estimator of coleopteran diversity in a wide arrange of environments. On the other hand, as an indicator of agricultural management, Silva et al. (2002) also found this family to be more diverse under a management that would fit into our “traditional” category.

In the agricultural context, curculionids are often thought of as a pest, because they feed on a wide range of crops (maize, beans, avocado, cotton, rice, etc.) (Morrone 2014). However, it is worth noting that none of the peasants interviewed in this study regarded them as an important problem (see Silva et al. 2002 for a similar case). In general, they mentioned that curculionids bit some of the stored grains, but the amount was not considered significant and these grains were generally used to feed fowl. Also, many of them said to avoid damage by storing the grains with local herbs and flowers which repel curculionids, while only a few of the more industrialized or market-oriented producers declared the use of industrial pesticides as part of their storage practices. In any case non of the interviewed peasants reported significant damage by curculionids in the field.

Finally, we have to warn about the limitations of this study. We only conducted one sampling (during the rainy season), which is restricted considering the high inter-annual, inter-geographic and inter-crop variability found in coleopterans (Finn et al. 1999; Andresen 2003). Hence, results must be taken cautiously. Also, our sample size was relatively small (16 plots), although the collaboration of key informants and the recollection of a large amount of beetles seemed to compensate for this, as shown by the clear differences between management types and beetle diversity (see Blanco & Castro 2007 for a discussion on sampling for qualitative information). Given that this was a field study, we had little control over environmental variables, which made statistical replicates impossible and limited the analysis possibilities. As a counterpart, those results that were statistically significant even under the great amount of environmental noise are probably a reflection of tendencies that are stronger than we could observe.

In this study we were able to organize the large heterogeneity of agricultural managements in Zaachila, Oaxaca, into two types. From these types, the one characterized by the use of local landraces and little external inputs exhibited more coleopteran diversity than its more industrialized counterpart. In particular, curculionids maintained a good correlation with overall coleopteran biodiversity, as well as with management intensification. In another work in Zaachila (Urrutia et al. 2019), we showed that agricultural patches, specially rainfed-agriculture patches, dominate this rural landscape and that, given their atomized pattern and large connectivity, they play a central role in the potential migration and recolonization among the adjacent forest patches. Together, these two studies suggest that traditional agriculture being practiced in the agricultural patches of Zaachila would render a high quality matrix for biodiversity conservation. These and other works support a key role of traditional *campesino* agriculture in the conservation of biodiversity, including agrobiodiversity and its cultural expressions (Perfecto et al. 2009; Mora 2017; Bellon et al. 2018) in this type of peasant-driven landscape.

## Acknowledgments

The authors thank members of *La Parcela* Laboratory for their valuable comments and suggestions. Raymundo Aguilar, Alexandre Beaupré, Maria de Guadalupe León, members of *El Molote* collective and the *Regiduría de agricultura del Municipio de Villa de Zaachila* helped in the field trips and recognition of the plots. Cecilia González González acknowledges the graduate program “Posgrado en Ciencias Biológicas, Universidad Nacional Autónoma de México” and CONACyT scholarship M. Benítez acknowledges financial support from UNAM-DGAPA-PAPIIT (IN207819). L. Jardón Barbolla acknowledges financial support form UNAM-DGAPA-PAPIIT (IA202515) and is grateful for NOT being part of the Sistema Nacional de Investigadores. MB, LJB, CGG and TLG conceived the ideas and designed methodology; CGG and TLG collected the data; CGG analysed the data; CGG and MB led the writing of the manuscript. All authors contributed critically to the drafts and gave final approval for publication.

## Footnotes

1 The equation that relates *alpha* and *H-alpha* is *H-alpha=1/(1-alpha)*\**log(sum(pi*^*alpha))*. For more details, look at the work by Tóthmérész (1995) and Jost (2006).

## Notes

https://github.com/laparcela/Coleoptera

