## Supplemental Material for "Linking coleopteran diversity with agricultural management of maize agroecosystems in Oaxaca, Mexico"

**Supplementary material**

Figure S1. Location of the five sampling quadrants in each plot.

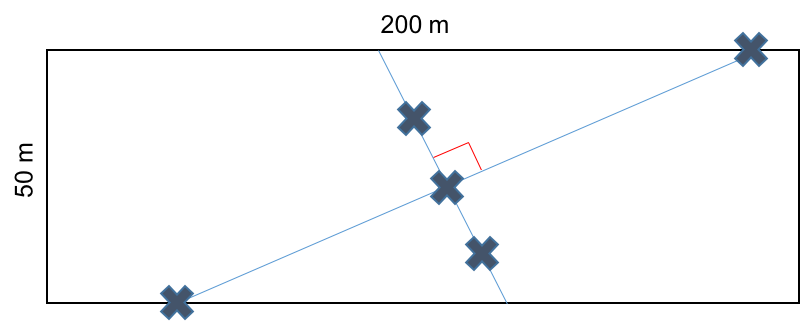

TableS2. List of al variables used for the typology construction, including their definition and data type.

| **Variable** | **Meaning** | **Data type** |
| --- | --- | --- |
| Number of crops | Total richness of planted species | counting |
| Number of varieties | Total richness of plant varieties taking all species into account | counting |
| Native seeds | Planting of native corn seeds | presence/absence |
| Monoculture | Planting scheme in which a single species is actively grown inside the plot | presence/absence |
| Rotation | Temporal or spatial rotation of crops in the plot | presence/absence |
| Fallow period | The habit of leaving land unused for at least one planting season | presence/absence |
| Animal traction | Use of animals at some part of the land preparation process, as opposed to tractors | presence/absence |
| Irrigation | Availability of irrigation water at any part of the year | presence/absence |
| Quelites | Habit of allowing useful weeds to grow inside the plots | presence/absence |
| Managed border | Habit of actively growing certain plants at the plot's borders | presence/absence |
| Trees | Presence of trees inside the plot or at its borders | presence/absence |
| Green manure | Habit of leaving uprooted or sown plant parts that serve as soil amendment and are grown specifically for this purpose | presence/absence |
| Stubble | Crop residues that are left in the field after harvesting | presence/absence |
| Compost | Any type of compost application to the field | presence/absence |
| Manure | Animal feces left in the field for purposes of soil amendment | presence/absence |
| Subsistence agriculture | Use of any part of the harvest for family consumption | presence/absence |
| Commercial agriculture | Use of any part of the harvest for market sell at any scale | presence/absence |
| Industrial fertilizer | Use of any commercial non-organic fertilizer | presence/absence |
| Industrial pesticide | Use of any commercial non-organic pesticide | presence/absence |
| Industrial herbicide | Use of any commercial non-organic herbicide | presence/absence |
| Organic fertilizer | Use of any commercial organic fertilizer | presence/absence |
| Organic pesticide | Use of any commercial organic pesticide | presence/absence |

Table S3. Table of values used for the FAMD analysis (the “crop names” and “variety names” columns are merely explanatory and were not introduced in the analysis).

| Plot | Crop names | Variety names | Number crops | Number varieties | native seeds | monoculture |
| --- | --- | --- | --- | --- | --- | --- |
| 1 | Maize, squash | White native maize; *tamala, chompa* and *huiche* squash | 2 | 4 | 1 | 0 |
| 2 | Maize, squash | Hybrid maize, *chompa* squash | 2 | 2 | 0 | 0 |
| 3 | Maize, squash, lemon, walnut | White native maize; *tamala, chompa* and *huiche* squash; native lemon, native walnut | 4 | 6 | 1 | 0 |
| 4 | Maize | *San José* hybrid maize | 1 | 1 | 0 | 1 |
| 5 | Alfalfa (*Medicago sativa*) | Commercial alfalfa | 1 | 1 | 0 | 1 |
| 6 | Maize, squash, beans | White native maize; *tamala, chompa* and *huiche* squash; native beans | 3 | 5 | 1 | 0 |
| 7 | Maize | White native maize | 1 | 1 | 1 | 1 |
| 8 | Maize | White native maize | 1 | 1 | 1 | 1 |
| 9 | Maize | Hybrid A7573 maize | 1 | 1 | 0 | 1 |
| 10 | Maize | White native maize | 1 | 1 | 1 | 1 |
| 11 | Maize, squash, alfalfa, lemon, chili pepper, nopal (*Opuntia*), beans | Yellow native maize, *huiche* squash, native beans, commercial alfalfa, *solterito* chili pepper, lemon, nopal | 7 | 7 | 1 | 0 |
| 12 | Maize, squash | White native maize; *tamala, chompa* and *huiche* squash | 2 | 4 | 1 | 0 |
| 13 | Maize | White native maize | 1 | 1 | 1 | 1 |
| 14 | Maize, beans, squash | White native maize; black native beans; *tamala, chompa* and *huiche* squash | 3 | 5 | 1 | 0 |
| 15 | Alfalfa, nopal | Commercial alfalfa, nopal | 2 | 2 | 0 | 1 |
| 16 | Maize | *Artillero* hybrid maize | 1 | 1 | 0 |  |

| Plot | rotation | fallow period | animal traction | irrigation | quelites | managed border | trees | green manure | stubble |
| --- | --- | --- | --- | --- | --- | --- | --- | --- | --- |
| 1 | 0 | 1 | 0 | 0 | 1 | 0 | 0 | 0 | 1 |
| 2 | 0 | 1 | 0 | 0 | 0 | 0 | 0 | 0 | 0 |
| 3 | 0 | 1 | 0 | 0 | 1 | 0 | 1 | 1 | 1 |
| 4 | 1 | 1 | 0 | 0 | 0 | 0 | 0 | 0 | 1 |
| 5 | 1 | 0 | 0 | 1 | 0 | 0 | 0 | 0 | 1 |
| 6 | 1 | 1 | 1 | 0 | 1 | 1 | 0 | 0 | 1 |
| 7 | 1 | 0 | 0 | 1 | 0 | 0 | 1 | 0 | 0 |
| 8 | 0 | 1 | 0 | 0 | 0 | 0 | 0 | 0 | 0 |
| 9 | 1 | 0 | 0 | 1 | 0 | 0 | 0 | 0 | 0 |
| 10 | 1 | 1 | 0 | 1 | 1 | 0 | 0 | 0 | 1 |
| 11 | 1 | 0 | 1 | 1 | 1 | 1 | 1 | 0 | 0 |
| 12 | 1 | 0 | 1 | 1 | 1 | 0 | 1 | 0 | 0 |
| 13 | 0 | 0 | 0 | 1 | 0 | 0 | 1 | 0 | 1 |
| 14 | 1 | 1 | 0 | 0 | 1 | 0 | 1 | 1 | 1 |
| 15 | 1 | 0 | 0 | 1 | 1 | 1 | 1 | 1 | 1 |
| 16 | 1 | 0 | 0 | 1 | 1 | 0 | 0 | 0 | 1 |

| Plot | compost | manure | subsistence agriculture | commercial agriculture | industrial fertilizer | industrial pesticide | industrial herbicide | organic fertilizer | organic pesticide |
| --- | --- | --- | --- | --- | --- | --- | --- | --- | --- |
| 1 | 0 | 0 | 1 | 1 | 0 | 0 | 0 | 0 | 0 |
| 2 | 0 | 1 | 1 | 1 | 1 | 1 | 0 | 0 | 0 |
| 3 | 0 | 1 | 1 | 1 | 1 | 0 | 1 | 1 | 0 |
| 4 | 0 | 0 | 0 | 1 | 1 | 0 | 1 | 1 | 0 |
| 5 | 0 | 1 | 0 | 1 | 1 | 1 | 1 | 0 | 0 |
| 6 | 0 | 0 | 1 | 1 | 1 | 0 | 0 | 0 | 0 |
| 7 | 0 | 1 | 1 | 0 | 0 | 1 | 0 | 0 | 0 |
| 8 | 0 | 1 | 1 | 1 | 1 | 0 | 1 | 0 | 0 |
| 9 | 0 | 0 | 0 | 1 | 1 | 1 | 0 | 0 | 0 |
| 10 | 0 | 0 | 1 | 1 | 1 | 0 | 1 | 0 | 0 |
| 11 | 1 | 1 | 1 | 1 | 0 | 1 | 0 | 1 | 0 |
| 12 | 0 | 1 | 1 | 1 | 0 | 1 | 0 | 0 | 0 |
| 13 | 0 | 0 | 0 | 1 | 1 | 0 | 1 | 1 | 0 |
| 14 | 0 | 0 | 1 | 1 | 0 | 0 | 0 | 1 | 0 |
| 15 | 1 | 1 | 0 | 1 | 0 | 0 | 0 | 1 | 1 |
| 16 | 0 | 0 | 1 | 0 | 1 | 1 | 1 | 0 | 0 |
